# Genome Evolution of *Acinetobacter baylyi* ADP1 During Laboratory Domestication: Acquired Mutations Impact Competence and Metabolism

**DOI:** 10.1101/2025.05.05.652239

**Authors:** Isaac Gifford, Meghna R Vergis, Jeffrey E Barrick

## Abstract

The bacterium *Acinetobacter baylyi* is a model organism known for its extreme natural competence and metabolic versatility. It is capable of transforming environmental DNA at a high frequency across all growth phases. The type strain ADP1 was created by random mutagenesis of a precursor strain, BD4, to prevent it from forming cell chains in culture. ADP1 has since been distributed between research groups over several decades and acquired subsequent mutations during this time. In this study we compare the genome sequences of *Acinetobacter baylyi* BD4 and its modern descendants to identify and understand the effects of mutations acquired and engineered during its domestication. We demonstrate that ADP1 variants in use today differ in their competence, growth on different carbon sources, and autoaggregation. Additionally, we link the global carbon storage regulator CsrA and a transposon insertion that removes its C-terminal domain specifically to changes in both overall competence and an almost complete loss of competence during stationary phase. Reconstructing the history of ADP1 and the diversity that has evolved in the variants currently in use improves our understanding of the desirable properties of this experimentally and industrially important bacterium and suggests ways that its reliability can be improved through further genome engineering.

**Significance:** *Acinetobacter baylyi* ADP1 is a bacterial chassis of interest to microbiologists in academia and industry due to its extreme natural competence and wide metabolic range. Its ability to take up DNA from its environment makes it straightforward to efficiently edit its chromosome. We identify and characterize mutations that have been passed down to modern strains of ADP1 from the initial work on ADP1 in the 1960s as well as subsequent mutations and genome edits separating strains in use by different research groups today. These mutations, including ones in a global regulator, have significant phenotypic consequences that have affected the reproducibility and consistency of experiments reported in the literature. We link a mutation in this global regulator to unexpected changes in natural competence. We also show that domesticated *A. baylyi* strains have impaired growth on a variety of carbon sources.

## Introduction

Microbes isolated from natural environments frequently find uses in laboratories as model systems for understanding biology or platforms for industrial applications. Many of these strains have been intentionally mutated to improve specific properties [1,2]. Even when strains have not been intentionally engineered, however, their genotypes do not remain static over time due to evolution [3–7]. Because isolates of commonly used strains are also frequently shared between research groups, mutants with altered properties can also be unintentionally disseminated. Mutants may lose traits unnecessary in laboratory cultures, such as motility [8] or extraneous metabolic pathways [5,7]. These changes in phenotype are frequently the result of mutations in global regulators that control several genes with varied functions [3,7]. Growth of strains in lab cultures can also unintentionally result in the gain of phenotypes such as antibiotic resistance [4]. This evolution can occur on short timescales, due to a single mutation taking over within a population after a few generations of use in the lab [3,4], or on longer timescales as multiple mutations accumulate over years of routine use [2,4].

The bacterium *Acinetobacter baylyi* ADP1 has been used in a broad range of applications in academic and industrial settings [9]. ADP1 is highly naturally competent, able to bind and take up DNA in a non-sequence specific manner from the environment through the action of its competence pilus [10]. It is known for the rare and perhaps unique trait of maintaining high competence in culture across different stages of normal growth [11,12]. This trait makes it straightforward to engineer its genome [13], as well as making it a model organism for studying the mechanism of natural competence [10] and its impact on genome evolution [14]. ADP1 is also of interest for industrial applications, ranging from lignin valorization to bioremediation, because it can consume and synthesize a variety of relevant and desirable compounds [15–17].

The BD4 strain that would become ADP1 was originally isolated from soil based on its ability to grow on 2,3-butandiol [18]. Subsequently, Elliot Juni and Alice Janik mutated the strain with UV to isolate a colony that did not form cell chains in culture and had a reduced capsule [1]. This strain was designated BD413. Elliot Juni deposited these original strains, BD4 and BD413, with the American Type Culture Collection under the accession numbers ATCC 33304 and ATCC 33305, respectively. These strains were originally classified as *Acinetobacter calcoaceticus*, then as an unnamed *Acinetobacter* species, and finally as *Acinetobacter baylyi* [19]. Strain BD413 has subsequently been passed between labs over the last six decades. The ADP1 designation of a BD413 stock from Nick Ornston’s lab is the standard one used for this strain today [19,20].

In this study we investigate mutations that evolved in *A. baylyi* ADP1 over six decades of laboratory domestication. We identify mutations tied to the original UV mutagenesis and laboratory propagation that resulted in multiple descendents of BD413 that are all referred to in the literature as ADP1. These ADP1 variants and the transposon-free variant ADP1-ISx (“ISx”) that was engineered from one of them [21] bear marked genotypic and phenotypic differences that affect key *A. baylyi* features including its competence and growth on a variety of carbon sources. These findings will aid in the design of future experiments utilizing this model organism and promote clarity and reproducibility between different research groups.

## RESULTS

### *A. baylyi* lost a 130-kb plasmid during domestication

We constructed a reference genome of *A. baylyi* strain ATCC 33304, identified as the ancestral strain BD4, from Nanopore reads and polished the assembly with Illumina reads (**Supplementary File 1**). It consists of two circular contigs of 3,648,419 and 130,836 base pairs. These were annotated with Prokka [22] and ISEscan [23] to identify coding regions and insertion elements. Prokka identified 3314 protein coding genes, 76 tRNAs, and seven sets of rRNA genes in the chromosomal assembly. ISEscan identified two complete IS elements in the chromosome: one IS*3* and one IS*256*. The smaller contig resembles a plasmid, and was designated pBD4-1. This plasmid has 120 protein coding genes and belongs to the R3-T7 group of *Acinetobacter* plasmids [24]. It contains a p*dif* module [25] consisting of two p*dif* sites located approximately 5.3 kb apart. pBD4-1 contains twelve complete IS elements that are predicted to be members of the IS*3*, IS*5*, IS*982*, and ISNCY families. The pBD4-1 plasmid was not present in any of the derived *A. baylyi* strains (**Fig. 1, Supplementary Table 3**), suggesting it was lost early in domestication.

**Figure 1.**
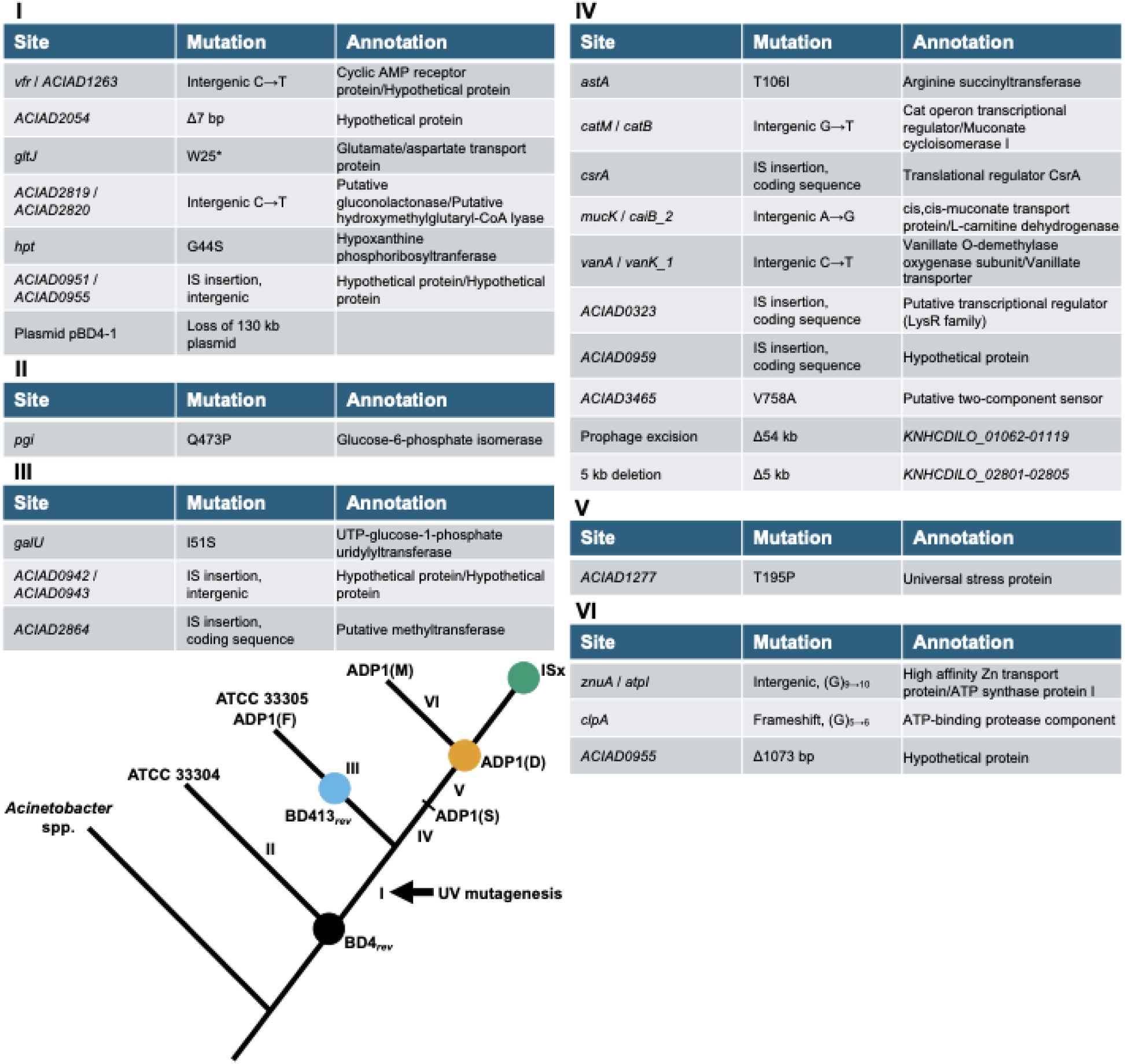
Phylogeny of *A. baylyi* strains inferred from mutations that evolved during domestication. Branch labels correspond to sets of mutations that evolved along each branch. Targeted deletions of IS elements during the creation of strain ISx are described in [21]. Circles indicate strains used in assays in this study.

### Domesticated *A. baylyi* strains diverged into two major clades

Illumina sequencing reads from six domesticated *A. baylyi* strains (**Supplementary Table 1**) were compared against the reference genome assembled for ATCC 33304. A point mutation in *pgi* was predicted in all other strains. However, our subsequent analysis demonstrated that this reflects a mutation that evolved in the ATCC 33304 strain and was not present in the original BD4 ancestor (see “Domestication of ADP1 altered cell morphology,” below) rather than a mutation arising in the domesticated lineage. Extant strains descended from *A. baylyi* BD4 differed from ATCC 33304 by as many as twenty-one mutations (**Fig. 1**). We inferred a maximum parsimony phylogenetic tree from the presence and absence of mutations in each genome. Seven mutations were present in all domesticated strains (**Fig. 1**, group I). Presumably, these occurred during or soon after mutagenesis by Juni and Janik. The domesticated strains then diverged into two groups depending on the presence or absence of either two specific IS insertions (**Fig. 1**, group III) or an additional ten mutations (**Fig. 1**, group IV). The first clade includes ATCC 33305, deposited at ATCC by Juni. It likely represents minimal adaptation to laboratory culture since the original mutagenesis of BD4. This strain was identical to the one used by the Friedman Lab, designated ADP1(F) here [26]. The second clade is composed of three other strains used by present-day research groups, and thus the ten additional mutations they share arose during routine use before a stock was distributed to these labs. The ADP1(D) and ADP1(M) [14] strains share a point mutation in a universal stress protein (**Fig. 1**, group V) not found in the ADP1(S) [27] strain. The ADP1(M) strain also acquired an additional three mutations not found in the other two (**Fig. 1**, group VI).

### Domestication of ADP1 altered cell morphology

Strains ATCC 33304 and 33305 grew in large aggregates in LB media (**Fig. 2A**) whereas ADP1 and ISx grew as single cells. ATCC 33304 and 33305 each contained a mutation in a gene related to capsule biosynthesis: either *pgi* or *galU*, respectively (**Fig. 1**, groups II and III, and **Fig. 2B**). We reverted these mutations to create strains BD4*_rev_*and BD413*_rev_* (**Fig. 2C**), which are expected to be genetically identical or nearly so to the original BD4 and BD413 strains [1]. As BD4 did not form aggregates in Juni and Janik’s original paper [1] this confirms that the *pgi* mutation arose later, as depicted in **Figure 1**. Both BD4*_rev_* and BD413*_rev_*grew as single cells in LB media and in the S2 minimal glucose media used by Juni and Janik [1] (**Fig. 3A**), consistent with the results of their original mutagenesis. However, BD4*_rev_*, BD413*_rev_*, and ADP1 all grew as cell chains when succinate, commonly used as a carbon source for ADP1, was substituted for glucose (**Fig. 3B**). All three strains, including BD4*_rev_*, grew as single cells in LB medium (**Fig. 3C**).

**Figure 2.**
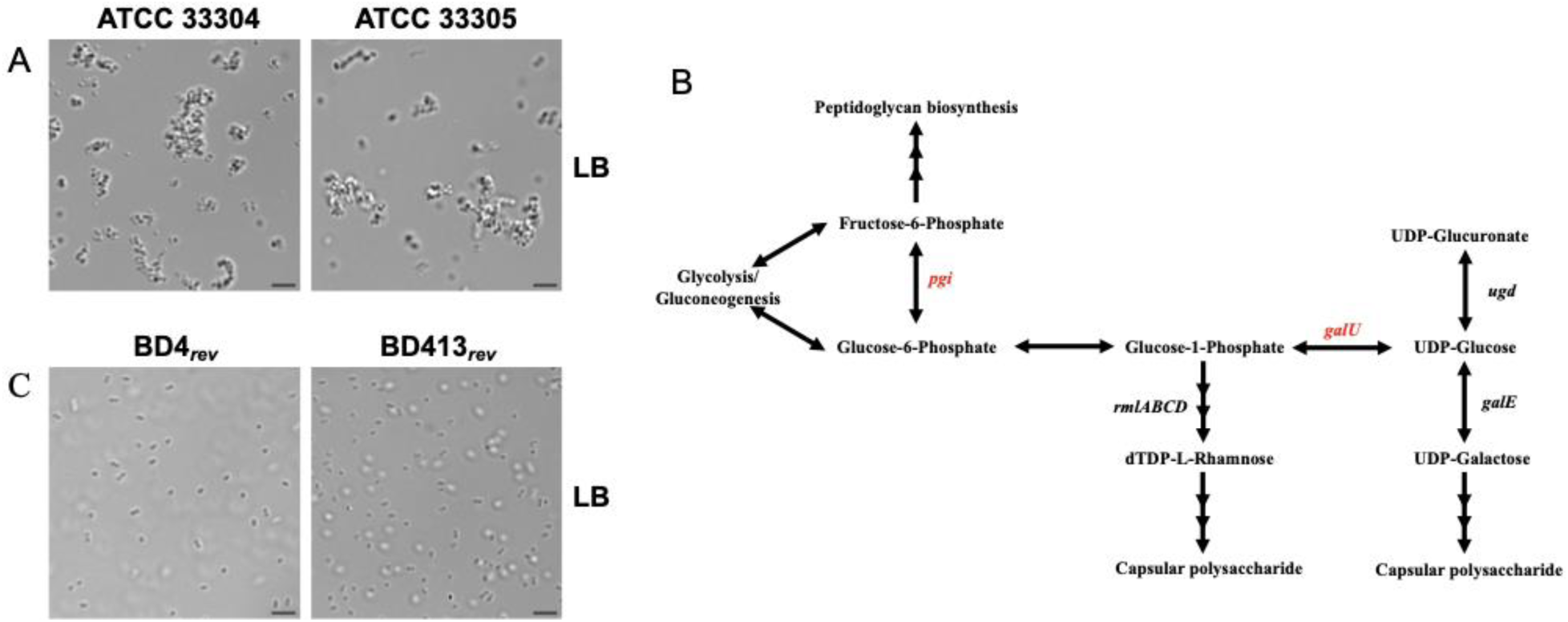
Mutations leading to aggregation in *A. baylyi* strains. (A) Microscope images of strains ATCC 33304 and 33305 grown in LB medium. (B) Metabolic pathways showing conversion of glucose to capsular polysaccharides. Genes mutated in ATCC 33304 and 33305 are shown. Pathways adapted from KEGG [28]. (C) Reverting *pgi* or *galU* mutations in ATCC 33304 and 33305, respectively, produced single cells in LB. Scale bars represent 5 μM.

**Figure 3.**
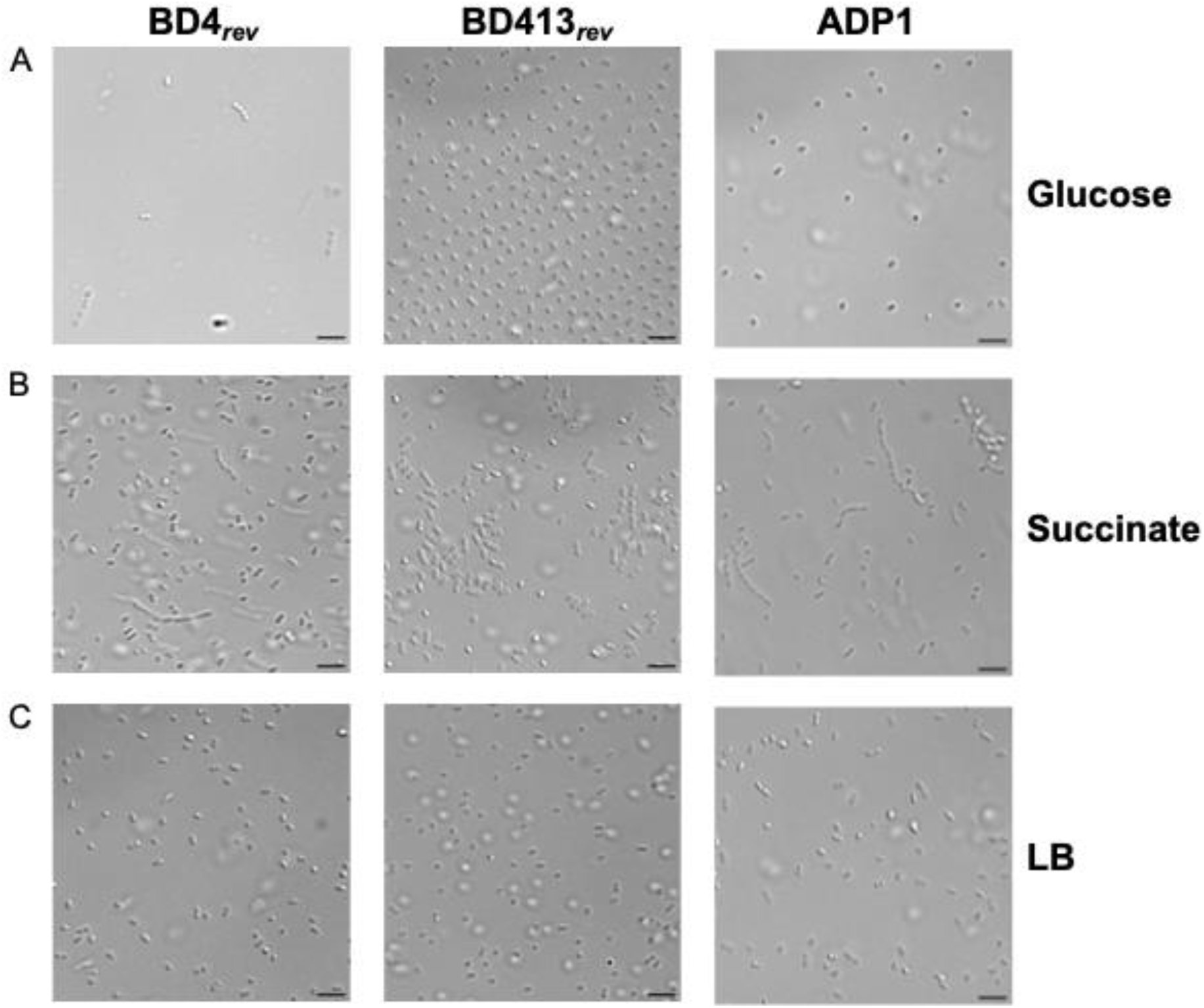
*A. baylyi* strains form cell chains in minimal media. (A) Domestication in BD413*_rev_* and ADP1 prevented cell chains from forming in S2 medium with glucose. (B) Cell chains formed for all strains when succinate was used as a carbon source in S2 medium. (C) All strains, including BD4*_rev_*, grew unicellularly in LB medium. Scale bars are 5 μM.

### Domestication of ADP1 affected competence

Previous reports in the literature have differed on the competence of BD4 [1] and BD413 [11], particularly in their competence during stationary phase. To investigate whether such differences may have been caused by domestication we performed competence assays on BD4*_rev_*, BD413*_rev_*, ADP1, and ISx. The domesticated strain ADP1 was significantly less competent than BD4*_rev_*, BD413*_rev_*, and ISx (pairwise Welch’s *t*-tests, adj. *p* = 0.0025, 0.019, and 0.027, respectively). About fifteen-fold fewer ADP1 cells incorporated an antibiotic resistance marker with flanking homology to the chromosome in an overnight transformation assay (**Fig. 4A**). Competence in the other strains did not vary significantly. We sought to investigate whether the difference in competence in ADP1 was caused by changes in competence during different growth phases (**Fig. 4B**). BD4*_rev_* and BD413*_rev_* maintained high levels of competence in both log (three hours post-inoculation) and stationary (twenty-four hours post-inoculation) phases. ADP1, however, had >2000-fold reduced competence during stationary phase. A similar loss was not observed for ADP1-ISx (“ISx”), the engineered derivative of ADP1. For it, we observed a 10-fold decrease in transformation efficiency during stationary phase, but this difference was not statistically significant (adj. *p* = 0.4). These results suggest that the reduced competence of ADP1 is due to a transposon insertion that truncated the global regulator *csrA* during domestication, a mutation that is unique to ADP1 and was removed during the construction of ISx (**Fig. 1**). A strain of ISx with the truncation recreated exhibited a similar >1000-fold decrease in competence during stationary phase (**Fig. 4C**).

**Figure 4.**
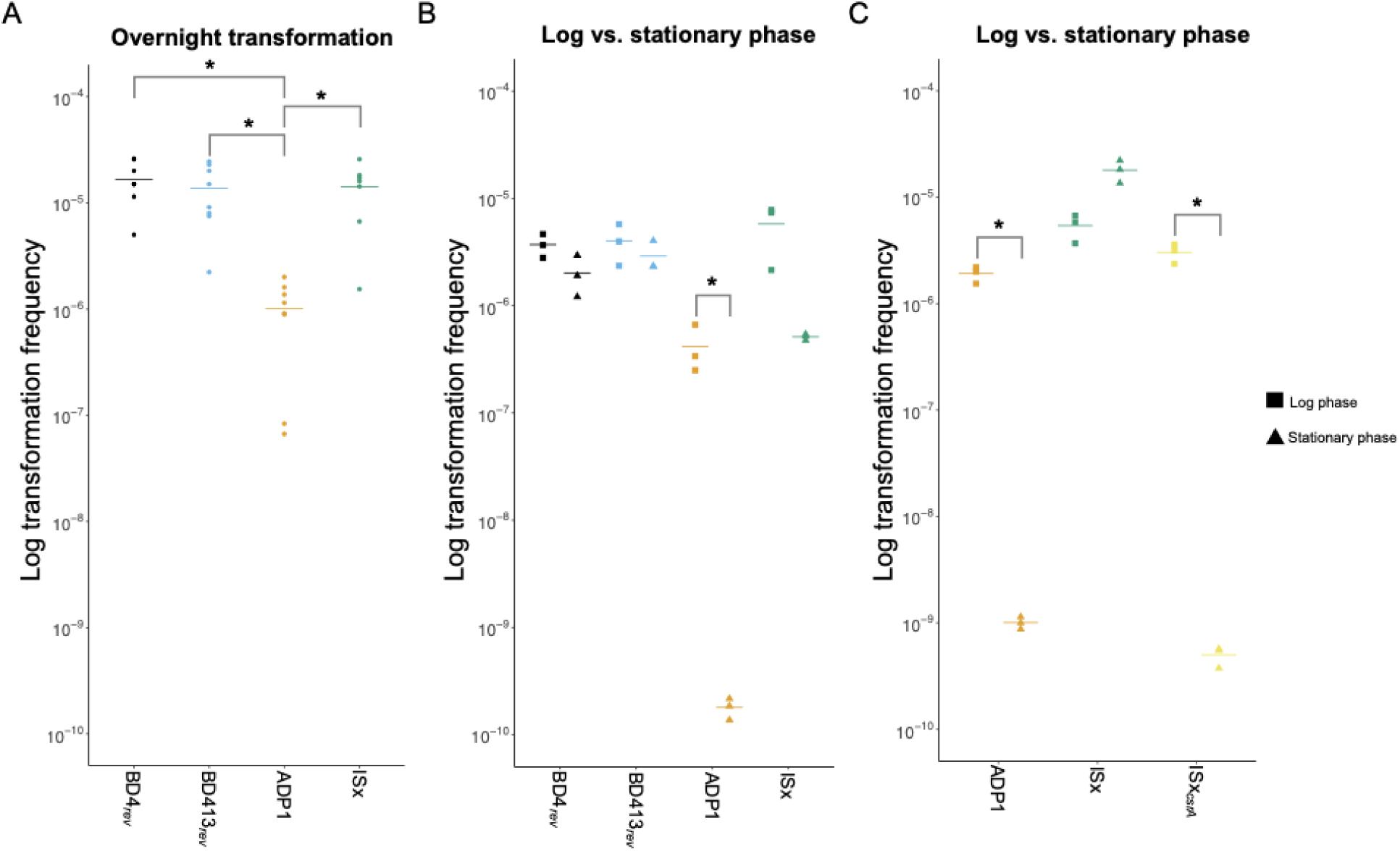
Transformation efficiency varies between domesticated *A. baylyi* strains. (A) Transformation frequencies of BD4_rev_ (black), BD413_rev_ (blue), ADP1 (orange), and ISx (green) strains incubated overnight with genomic DNA carrying a *specR* resistance gene. (B) Transformation frequencies of the same strains during log (squares) and stationary (triangles) phases. At each time point, each strain was incubated with transforming DNA for 30 minutes, then the remaining free DNA was digested by DNase I before plating to count transformants. (C) Transformation frequencies during log (squares) and stationary (triangles) phases for ADP1 and ISx compared to an ISx mutant with a truncated *csrA* gene (yellow). \**p* < 0.05 for Welch’s *t*-tests.

### Domestication of ADP1 impacted growth across different carbon sources

We investigated growth of domesticated *A. baylyi* strains on different carbon sources to determine whether strain choice impacts its role as a model system for the degradation of aromatic compounds or as a chassis for metabolic engineering. When grown in LB media, ADP1 and ISx grew to a substantially higher cell density than BD4*_rev_* and BD413*_rev_*but did not show significantly higher growth rates (**Fig. 5A**). Further experiments were carried out in modified S2 media as described by Juni and Janik [1] but with other carbon sources substituted for glucose. On glucose BD4*_rev_* reached its maximum density substantially faster than the domesticated strains, with ADP1 lagging further behind BD413*_rev_* and ISx, whereas on succinate (160 g/l) there was no noticeable difference between the four strains (**Fig. 5BC**). The growth defect in the domesticated strains on glucose was rescued by adding a small amount of succinate to the media (16 g/l, 1/10^th^ the normal concentration) (**Fig. 5D**). However, BD4*_rev_* displayed a two-step growth curve typical of catabolite repression in the mixed media whereas the other strains showed only a single log phase. When grown on the aromatic compounds benzoate (0.24 g/l) and vanillate (3.4 g/l), the laboratory domesticated strains ADP1 and ISx grew slower than BD4*_rev_* or BD413*_rev_* (**Fig. 5EF**). ADP1 grew to a particularly low optical density on vanillate, and truncating *csrA* in the way that evolved in ADP1 reconstituted this phenotype in ISx.

**Figure 5.**
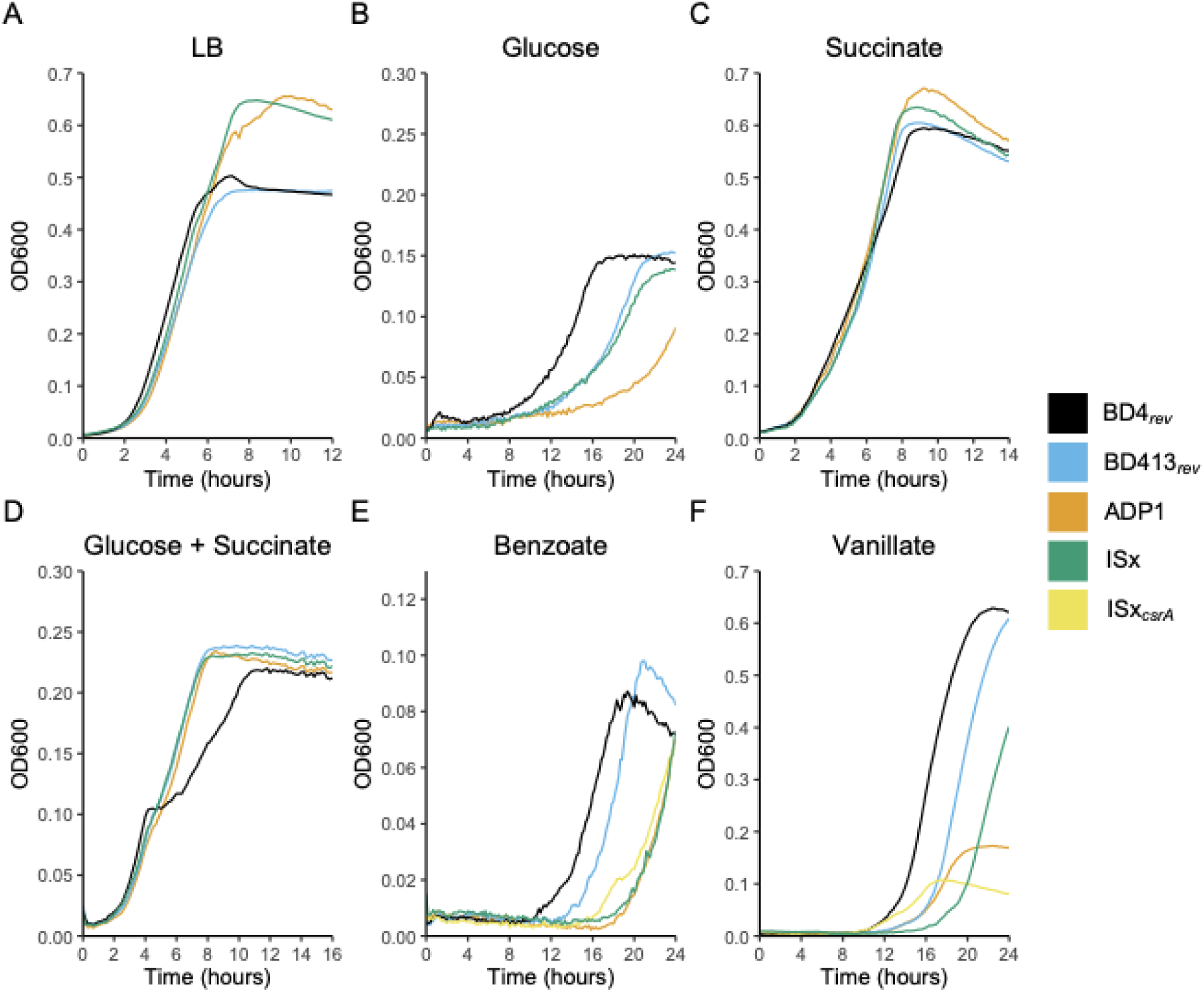
Domestication impacted growth and catabolite repression in *A. baylyi*. Growth curves of *A. baylyi* strains in (A) LB medium and in S2 media with (B) 3 mM glucose, (C) 25 mM succinate, (D) 3 mM glucose and 2.5 mM succinate, (E) 2 mM benzoate, or (F) 20 mM vanillate. Lines represent the average OD_600_ of six replicates each. The strains tested were BD4*_rev_* (black), BD413*_rev_* (blue), ADP1 (orange), and ISx (green). E and F include an ISx mutant with the *csrA* truncation from ADP1 (yellow).

### Adaptation of BD4_rev_ to laboratory culture

To investigate which mutations that evolved in ADP1 were likely caused by adaptation to laboratory culture we “replayed” its domestication by evolving a BD4*_rev_* strain in minimal succinate media (ABMS). At the end of one month of daily serial transfers (∼300 bacterial generations), we sequenced the genomes of endpoint clonal isolates, one each from eleven separate populations. Each isolate evolved between zero and three mutations, averaging two mutations per genome (**Fig. 6A**). The predominant gene mutated was *ACIAD1238*, a stress response regulator. It experienced IS*982* element insertions in six isolates at four different positions between the −10 and −21 positions upstream of its start codon. The second most commonly mutated gene was Ribonuclease D (*rnd*), which was disrupted in five endpoint isolates. Mutations in *rnd* included one out-of-frame insertion, one nonsense mutation, two missense mutations, and an IS*3* element insertion. One isolate that did not have a mutation in *rnd* had a missense mutation in the coding sequence of *hfq*, an RNA chaperone. Two isolates that had mutations in *rnd* also had base substitutions either 88 bp or 90 bp upstream of *hfq*. One isolate that did not have a mutation in *rnd* had a substitution in the −7 position upstream of *csrA*. One isolate had a 36-bp deletion in the middle of *ACIAD1854*, a putative phage-related protein. An IS*982* element transposed from the BD4*_rev_* plasmid into the chromosome 234 bp upstream of *ACIAD2521* and 126 bp downstream of *ACIAD2522* in three clonal isolates.

**Figure 6.**
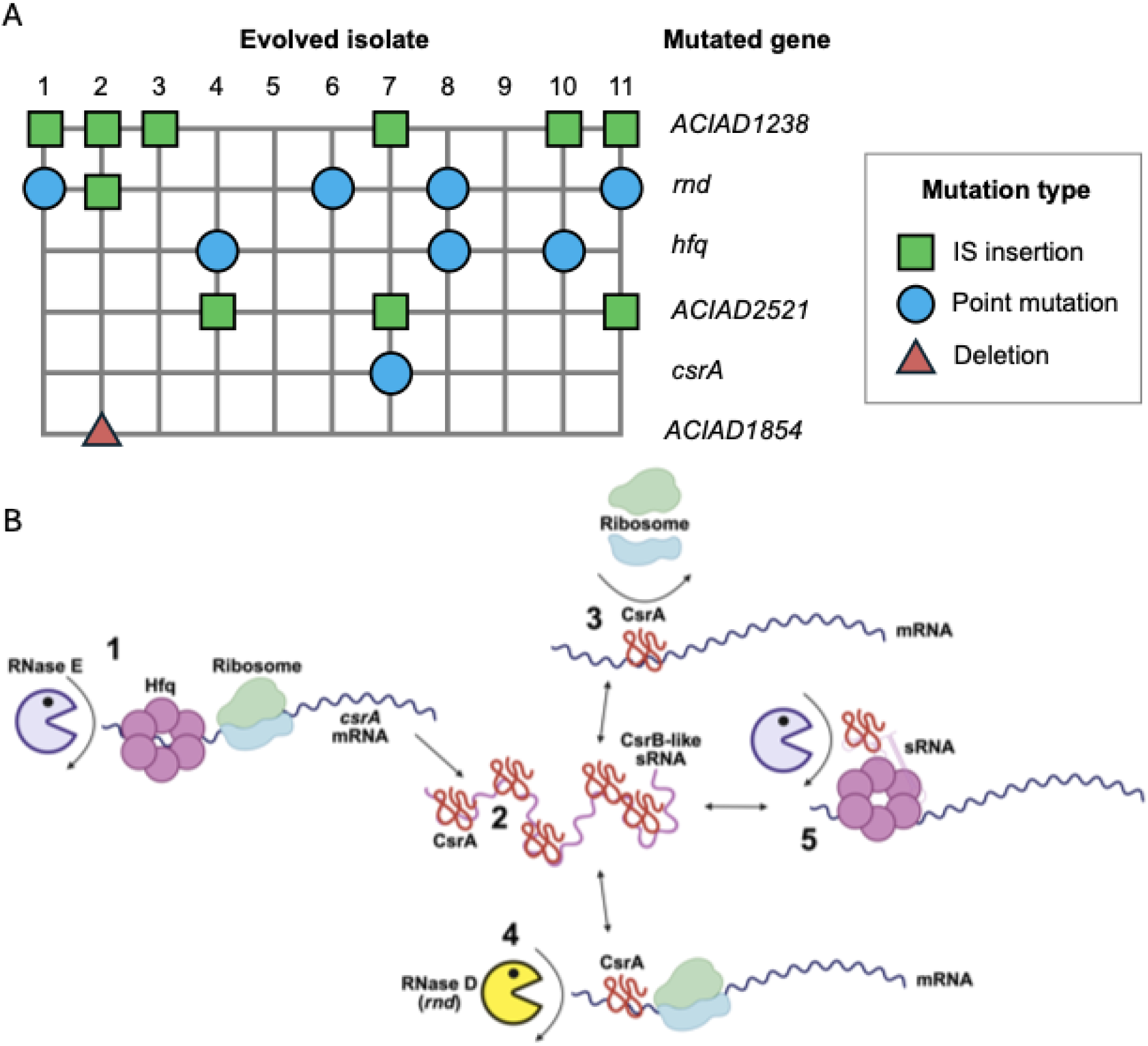
(A) Mutations evolved in BD4*_rev_* (MRV001) during one month of evolution in *A. baylyi* minimal succinate (ABMS) medium. (B) Potential post-transcriptional regulatory network inferred in *A. baylyi* from interactions between CsrA, Hfq, and RNase D. (1) Hfq protects *csrA* mRNA from degradation by RNase E [29]. (2) CsrA proteins are bound by a CsrB-like mRNA [30]. (3) CsrA binds to Shine-Dalgarno sites, preventing ribosome assembly on mRNAs [31]. (4) CsrA protects mRNAs from degradation by RNase D (*rnd*). (5) CsrA protects small regulatory RNAs from degradation by RNases [32]. Figure 6B created with Biorender.

## Discussion

Through comparative genomics we identified a set of mutations linked to the mutagenesis of *A. baylyi* BD4 that Juni and Janik performed to select for microencapsulation and unicellular growth, which have been inherited by its descendant strains. However, these mutations only appear to have had the desired effect in the S2 glucose medium: cell chains were still observed when the same strains were grown in the same base medium substituted with other carbon sources. Bacterial capsules can cause the formation of cell chains [33,34], thus it is likely that the microencapsulation and unicellular phenotypes are tied to one or a few of the mutations we have linked to the original mutagenesis. In this study we also demonstrated that some extant strains have acquired mutations causing autoaggregation. The capsule biosynthesis pathway has similarly mutated in ways that have caused auroaggregation in previous *A. baylyi* evolution experiments [35,36]. As bacterial capsules can block autoaggregation by masking cell surface proteins such as adhesins [37], it is likely that the complete loss of the capsule, caused by mutations during routine culturing, exposes these proteins and allows them to interact. Future work could further promote *A. baylyi*’s ease of use by engineering a permanent solution to the issues of cell chains and autoaggregation through targeted deletions of capsule biosynthesis and cell surface adhesion proteins. This solution would have the added benefit of rerouting metabolic flux away from capsule biosynthesis, which may improve yields of high-value products [38].

An unexpected finding was a novel 130 kb plasmid not found in any of the domesticated strains. This plasmid could have been lost during UV mutagenesis or routine passaging over time. It encodes several antibiotic and heavy metal resistance transporters, as well as metabolic enzymes for the catabolism of plant-derived compounds, suggesting that it had a role in bacterial survival and growth in the soil environment from which BD4 was originally isolated. During domestication, *A. baylyi* strains also evolved mutations in chromosomal copies of two genes with copies on this plasmid: *hpt* (hypoxanthine phosphoribosyltransferase) and *vanK* (vanillate transporter). These mutations may compensate for the loss of the plasmid-encoded copies.

We found that genetic diversity existed among several *A. baylyi* strains designated as ADP1. Divergence was especially notable between the strain deposited at ATCC (ATCC 33305) and strains from different research groups. ATCC 33305 has undergone minimal evolution since the original mutation performed by Juni and Janik [1]. Strains in the other major clade were all obtained from academic labs that inherited their strains from other labs. These strains have undergone further domestication and adaptation to laboratory culture over time. Strains in this clade evolved a shared set of mutations. Among these is the truncation of *csrA* [35], which led to a ten-fold decrease in transformation efficiency and an even larger decrease during stationary phase. Reversion of this mutation explains the increase in competence in the engineered ISx strain [21].

Reports on competence in *A. baylyi* have varied, particularly on its competence during different stages of growth. Juni and Janik originally described BD4 as a strain that was maximally competent at the onset of stationary phase, with competence decreasing during log and into stationary phases [1]. Subsequent work on BD413 described it as highly competent at all growth stages, including log phase [11,12]. We did not observe differences in competence between BD4*_rev_* and BD413*_rev_* or between log and stationary phase for either strain, supporting high competence across growth phases for *A. baylyi* strains with intact, ancestral *csrA* genes.

Previously we reported that the strain ISx, created by deleting all insertion sequences from ADP1, frequently adapted in evolution experiments through mutations in *rnd* (RNase D), *hfq*, and *csrA* [35], whereas ADP1 (with its truncated *csrA* gene) does not [36]. CsrA and Hfq are post-transcriptional regulators that affect the translation and stability of mRNAs (**Fig. 6B**). RNase D has also been shown in *Myxococcus xanthus* to regulate its lifestyle by degrading a specific transcript [39]. It is likely that interactions between these proteins form a post-transcriptional regulatory network in *A. baylyi*. Here, we found that these genes also mutated in BD4*_rev_* during an evolution experiment. This consistent result suggests that truncation of *csrA* evolved from general adaptation to laboratory culture during domestication, rather than in response to the mutations introduced by UV mutagenesis in the history of ADP1.

CsrA is canonically involved in the regulation of carbon storage and catabolism [40], thus its mutation in ADP1 may be linked to modification of these pathways in laboratory culture. ADP1 often also evolves reduced competence in laboratory experiments [36,41], which has been ascribed to multiple factors, including a fitness cost incurred by taking up homologous genomic DNA [41] and the emergence of a prophage that is thought to utilize its competence pilus as an attachment site [42]. Thus, it is also possible that the truncation of *csrA* was selected for during domestication for the reduction in competence or for its effect on multiple processes. In other bacteria, mutations in global transcriptional regulators that respond to starvation and stress, such as RpoS in *E. coli* [3,43] and *Salmonella enterica* [7], are common during laboratory domestication. Frequent mutation of the post-transcriptional regulatory network in experiments with *A. baylyi* suggests it may play an important role in these types of responses. During adaptive laboratory evolution BD4*_rev_* also frequently evolved mutations in a universal stress protein (**Fig. 6A**). Two domesticated ADP1 strains in our reconstructed phylogeny had point mutations in the coding region of a separate universal stress protein. These findings suggest that modulating stress responses in ADP1 adapts it to laboratory conditions. Whether its stress responses have become more or less active cannot be determined from these mutations alone.

Domesticated *A. baylyi* strains showed reduced or slower growth on glucose and the aromatic compounds benzoate and vanillate compared to BD4*_rev_*, which could reflect reduction or loss of extraneous metabolic pathways or trade-offs due to mutations that are beneficial for growth on a limited range of carbon sources used in laboratory cultures. These pathways are of particular interest to research groups for their applications in metabolic engineering [15,17,27], and these differences could affect the reproducibility of experiments and product yields. CsrA is classically involved in control of glycolysis and gluconeogenesis [40], and truncation of *csrA* slowed the growth of ADP1 on glucose relative to ISx. Truncation of *csrA* also impaired catabolism of vanillate. Additionally, mutations around the vanillate metabolism and transport genes *vanA* and *vanK* and the *cat* genes likely also contribute to the reduced growth of ADP1 and ISx on aromatic compounds.

In this study we have shown that strains of ADP1 being used in different research groups have substantially different phenotypes. We recommend researchers sequence their *A. baylyi* strains to determine exactly which genotype they are working with and include that information in their publications to improve reproducibility. Our results also suggest avenues for rational strain improvement through precision genome editing of ADP1 that can recapitulate the benefits of mutations that evolved during domestication (e.g., elimination of autoaggregation) without their side-effects and trade-offs (e.g., reduced competence and worse growth on glucose). Finally, reversion of mutations acquired during domestication may also improve the growth of *A. baylyi* strains on carbon sources such as aromatic compounds derived from lignin, an activity that is desirable for bioenergy and biomanufacturing applications.

## Methods

### Strains and culture conditions

Strains used in this study are described in **Supplementary Table 1**. *A. baylyi* strains were grown at 30° C with shaking at 200 RPM in an incubator for liquid cultures in either LB media or minimal media. The minimal medium (S2) was a modified version of that used by Juni and Janik (1969). *Acinetobacter* minimal succinate medium (ABMS) [44] was used for adaptive laboratory evolution. For growth assays and microscopy ferric chloride (0.01 g/l final concentration) was substituted for ferrous sulfate, as it substantially improved growth. Carbon sources were substituted for glucose as listed in **Figure 5**. Vanillate and benzoate stocks were prepared as described by Luo et al. [27]. When necessary for selection, kanamycin (Kan, 50 μg/ml) or azidothymidine (AZT, 200 μg/ml) was added to culture media.

### Genomic DNA extraction, sequencing, assembly, and analysis

For short-read sequencing on an Illumina instrument, genomic DNA was extracted with a PureLink Genomic DNA Kit (Invitrogen) and sequenced as reported previously [35]. For long-read sequencing, high molecular weight DNA was extracted from 1 ml of overnight culture grown in LB with a Quick-DNA HMW Magbead Kit (Zymo Research). DNA extracts were barcoded with an Oxford Nanopore Rapid Barcoding Kit 24 V14 and sequenced on a MinION Mk1C sequencer. To assemble the reference genome of ATCC 33304 (*A. baylyi* BD4) long reads were assembled using Trycycler [45] following initial assembly of subsampled reads with Flye [46]. The assembly was then polished with short reads using Polypolish [47]. A final polish was performed by identifying mutations from the same short read sequences by running *breseq* [48] and applying called mutations to the reference sequence. The assembled genome was then annotated with Prokka [22]. The AcinetobacterPlasmidTyping v3.0 database [24] was used to identify the rep type of plasmid pBD4-1. p*dif* modules were identified as described by Blackwell and Hall [24,25]

Whole-genome sequencing data from published studies of unmodified *A. baylyi* genomes [14,26,27] were downloaded from NCBI Sequence Read Archive for comparative analyses (accessions SRR15734327, SRR20766519, and SRR28120535). Mutations in each genome were identified by using *breseq* [48] to compare Illumina reads from each strain against the assembled genomes of ATCC 33304 and ADP1 [36].

### Microscopy

Cultures were imaged on a Zeiss Axiovert 200M fluorescence microscope in the UT Austin Microscope and Imaging Facility. Ten microliters of overnight culture were placed on a glass slide and covered with a coverslip then imaged using the 63× objective and DIC contrast. Afterwards images were cropped and contrast was adjusted by linear scaling in Fiji [49].

### Mutant construction

*A. baylyi* mutants were constructed using Golden Transformation [13] with the primers listed in **Supplementary Table 1**. Briefly, a *tdk*/*kan* selectable/counterselectable cassette was inserted into the chromosome following ligation to homologous flanks targeting the desired gene amplified by PCR and selected for insertion of plates containing kanamycin. Isolated colonies from these plates were then inoculated into cultures in LB-Kan medium and grown overnight. The intended mutation was then introduced by recombination with a rescue cassette containing the mutation and 1 kb homologous flanks on each end followed by selection for loss of the *tdk* gene on LB-AZT plates. Insertion of the *tdk*/*kan* cassette into *csrA* was unsuccessful, so for the construction of the *csrA* truncation mutant the cassette was inserted immediately downstream of the gene instead. Mutant constructs were verified by Illumina sequencing and comparison to the ADP1 reference genome with *breseq* as described above. When mutating the *pgi* or *galU* genes, isolated colonies were streaked onto an additional LB-Kan or LB-AZT plate and grown overnight to ensure isolation of pure clones.

### Competence assays

Competence was measured by transforming each strain with genomic DNA from an ISx derivative with a spectinomycin-resistance gene replacing gene *ACIAD2049*. There are no mutations in any of the strains near *ACIAD2049* that could affect the frequency of recombination. For measurements of overall competence, 500 ng of genomic DNA of the spectinomycin-resistant ISx strain was added to 35 μl of eight replicate overnight cultures of each strain in 500 μl of LB media and incubated overnight. The following day each transformation was serially diluted in ten-fold increments to 10^−8^ in sterile saline and 5 μl of the dilutions were spot plated on LB plates with- and without-spectinomycin and incubated overnight before counting colonies to calculate the transformation rates. Transformation frequencies were calculated by dividing the CFUs/ml determined from selective plates by the CFUs/ml determined from non-selective plates.

For measurements of competence during growth, overnight cultures of each strain were diluted 1:25 by adding 2 ml of culture to 48 ml of fresh medium in 250 ml flasks. At each time point (three hours for log phase and 24 hours for stationary phase) 500 μl aliquots were transferred to separate 25 ml culture tubes. These were inoculated with 500 ng of transforming genomic DNA and incubated for thirty minutes before adding 50 ng DNase I. Digestion of remaining DNA was allowed to proceed for ten minutes with continued incubation at 30° C before diluting each transformation and plating it on selective and non-selective plates as above.

### Growth assays

Growth curves were performed in a Tecan Infinite 200 Pro plate reader. For each medium, cultures were first inoculated into 5 ml cultures in 25 ml glass tubes and incubated overnight. 5 μl of these cultures were transferred to 5 ml of fresh medium and incubated overnight again to precondition. Then, 2 μl of each overnight culture was inoculated into 198 μl of medium in each well of a 96-well clear flat-bottom microplate (Corning Incorporated, catalog number: 3596). OD_600_ readings were taken in the plate reader every ten minutes for 24 hours for six biological replicates per strain per medium with incubation at 30° C and orbital shaking at an amplitude of 3.5 mm for seven minutes between measurements.

### Adaptive evolution of BD4*_rev_*

Eleven colonies of GFP-tagged BD4_rev_ (strain MRV001, *gfp* inserted in place of *ACIAD2049*) were inoculated into separate culture tubes containing ABMS medium [44]. These cultures were allowed to grow for 24 hours before being diluted 1000× into fresh ABMS medium. The cultures were transferred 30 times, which corresponds to ∼300 generations of evolution [35]. At this point, 1 ml of each culture was frozen in 20% (v/v) glycerol. One endpoint clonal isolate was obtained from each evolved population by streaking it onto an LB agar plate, then restreaking an isolated colony on another LB agar plate. The final isolated clones were picked and grown in LB media, and 1 ml of each resulting culture was frozen in 20% (v/v) glycerol. The genomes of these clones were sequenced as described above. A base substitution leading to a K107E mutation in *ACIAD3105*, which encodes a DUF4124 domain-containing protein, occurred in the ancestor MRV001 during strain construction. It was excluded from the analysis of evolved mutations.

## Acknowledgements

We thank Daniel Deatherage for assistance with genome sequencing. This work was supported by the Welch Foundation (F-1979), the National Science Foundation (MCB-2123996 and DEB-1951307), and The University of Texas at Austin College of Natural Sciences (Spark Grant).

## Data Availability

Genome sequencing data is available from the NCBI Sequence Read Archive (PRJNA1256545).

## Supplementary Tables and Figures

**Supplementary Table 1** *A. baylyi* strains used in this study.

**Supplementary Table 2** Primers used in this study.

**Supplementary Table 3** Mutations in domesticated *A. baylyi* strains identified by *breseq*.

**Supplementary File 1** Reference genome of *A. baylyi* strain ATCC 33304 (BD4) in GenBank format.

## References

1. Juni E, Janik A. Transformation of Acinetobacter calco-aceticus (Bacterium anitratum). J Bacteriol. 1969;98: 281–288.

2. Browning DF, Hobman JL, Busby SJW. Laboratory strains of Escherichia coli K-12: things are seldom what they seem. Microb Genom. 2023;9. doi:10.1099/mgen.0.000922

3. Eydallin G, Ryall B, Maharjan R, Ferenci T. The nature of laboratory domestication changes in freshly isolated Escherichia coli strains: Laboratory domestication ofE. coli. Environ Microbiol. 2014;16: 813–828.

4. Carroll SM, Xue KS, Marx CJ. Laboratory divergence of Methylobacterium extorquens AM1 through unintended domestication and past selection for antibiotic resistance. BMC Microbiol. 2014;14: 2.

5. Marks ME, Castro-Rojas CM, Teiling C, Du L, Kapatral V, Walunas TL, et al. The genetic basis of laboratory adaptation in Caulobacter crescentus. J Bacteriol. 2010;192: 3678–3688.

6. Artuso I, Lucidi M, Visaggio D, Capecchi G, Lugli GA, Ventura M, et al. Genome diversity of domesticated Acinetobacter baumannii ATCC 19606T strains. Microb Genom. 2022;8. doi:10.1099/mgen.0.000749

7. Eisenstark A. Genetic diversity among offspring from archived Salmonella enterica ssp. enterica serovar typhimurium (Demerec Collection): in search of survival strategies. Annu Rev Microbiol. 2010;64: 277–292.

8. Barreto HC, Cordeiro T, Henriques A, Gordo I. Rampant loss of social traits during domestication of a Bacillus subtilis natural isolate. Sci Rep. 2019;10. doi:10.1038/s41598-020-76017-1

9. Elliott KT, Neidle EL. Acinetobacter baylyi ADP1: transforming the choice of model organism. IUBMB Life. 2011;63: 1075–1080.

10. Leong CG, Bloomfield RA, Boyd CA, Dornbusch AJ, Lieber L, Liu F, et al. The role of core and accessory type IV pilus genes in natural transformation and twitching motility in the bacterium Acinetobacter baylyi. PLoS One. 2017;12: e0182139.

11. Palmen R, Vosman B, Buijsman P, Breek CK, Hellingwerf KJ. Physiological characterization of natural transformation in Acinetobacter calcoaceticus. J Gen Microbiol. 1993;139: 295–305.

12. Leong CG, Boyd CM, Roush KS, Tenente R, Lang KM, Lostroh CP. Succinate, iron chelation, and monovalent cations affect the transformation efficiency of Acinetobacter baylyi ATCC 33305 during growth in complex media. Can J Microbiol. 2017;63: 851–856.

13. Suárez GA, Dugan KR, Renda BA, Leonard SP, Gangavarapu LS, Barrick JE. Rapid and assured genetic engineering methods applied to Acinetobacter baylyi ADP1 genome streamlining. Nucleic Acids Res. 2020;48: 4585–4600.

14. Sezmis AL, Woods LC, Peleg AY, McDonald MJ. Horizontal gene transfer, fitness costs and mobility shape the spread of antibiotic resistance genes into experimental populations of Acinetobacter baylyi. Mol Biol Evol. 2023;40: msad028.

15. Arvay E, Biggs BW, Guerrero L, Jiang V, Tyo K. Engineering Acinetobacter baylyi ADP1 for mevalonate production from lignin-derived aromatic compounds. Metab Eng Commun. 2021;13: e00173.

16. Santala S, Santala V, Liu N, Stephanopoulos G. Partitioning metabolism between growth and product synthesis for coordinated production of wax esters in Acinetobacter baylyi ADP1. Biotechnol Bioeng. 2021;118: 2283–2292.

17. Salcedo-Vite K, Sigala J-C, Segura D, Gosset G, Martinez A. Acinetobacter baylyi ADP1 growth performance and lipid accumulation on different carbon sources. Appl Microbiol Biotechnol. 2019;103: 6217–6229.

18. Taylor WH, Juni E. Pathways for biosynthesis of a bacterial capsular polysaccharide i. J Bacteriol. 1961;81: 688–693.

19. Vaneechoutte M, Young DM, Ornston LN, De Baere T, Nemec A, Van Der Reijden T, et al. Naturally transformable Acinetobacter sp. strain ADP1 belongs to the newly described species Acinetobacter baylyi. Appl Environ Microbiol. 2006;72: 932–936.

20. Patel RN, Mazumdar S, Ornston LN. Beta-ketoadipate enol-lactone hydrolases I and II from Acinetobacter calcoaceticus. J Biol Chem. 1975;250: 6567–6567.

21. Suárez GA, Renda BA, Dasgupta A, Barrick JE. Reduced mutation rate and increased transformability of transposon-free Acinetobacter baylyi ADP1-ISx. Appl Environ Microbiol. 2017;83. doi:10.1128/AEM.01025-17

22. Seemann T. Prokka: rapid prokaryotic genome annotation. Bioinformatics. 2014;30: 2068–2069.

23. Xie Z, Tang H. ISEScan: automated identification of insertion sequence elements in prokaryotic genomes. Bioinformatics. 2017;33: 3340–3347.

24. Lam MMC, Koong J, Holt KE, Hall RM, Hamidian M. Detection and Typing of Plasmids in Acinetobacter baumannii Using rep Genes Encoding Replication Initiation Proteins. Microbiol Spectr. 2023;11: e0247822.

25. Blackwell GA, Hall RM. The tet39 determinant and the msrE-mphE genes in Acinetobacter plasmids are each part of discrete modules flanked by inversely oriented pdif (XerC-XerD) sites. Antimicrob Agents Chemother. 2017;61. doi:10.1128/AAC.00780-17

26. Meroz N, Livny T, Toledano G, Sorokin Y, Tovi N, Friedman J. Evolution in microbial microcosms is highly parallel, regardless of the presence of interacting species. Cell Syst. 2024;15: 930–940.e5.

27. Luo J, McIntyre EA, Bedore SR, Santala V, Neidle EL, Santala S. Characterization of highly ferulate-tolerant Acinetobacter baylyi ADP1 isolates by a rapid reverse engineering method. Appl Environ Microbiol. 2022;88: e0178021.

28. Kanehisa M, Goto S. KEGG: Kyoto Encyclopedia of genes and genomes. Nucleic acids research. 2000;28: 27–30.

29. McNealy TL, Forsbach-Birk V, Shi C, Marre R. The Hfq homolog in Legionella pneumophila demonstrates regulation by LetA and RpoS and interacts with the global regulator CsrA. J Bacteriol. 2005;187: 1527–1532.

30. Kulkarni PR, Cui X, Williams JW, Stevens AM, Kulkarni RV. Prediction of CsrA-regulating small RNAs in bacteria and their experimental verification in Vibrio fischeri. Nucleic Acids Res. 2006;34: 3361–3369.

31. Baker CS, Eöry LA, Yakhnin H, Mercante J, Romeo T, Babitzke P. CsrA inhibits translation initiation of Escherichia coli hfq by binding to a single site overlapping the Shine-Dalgarno sequence. J Bacteriol. 2007;189: 5472–5481.

32. Stenum TS, Holmqvist E. CsrA enters Hfq’s territory: Regulation of a base-pairing small RNA. Mol Microbiol. 2022;117: 4–9.

33. Nachtigall C, Vogel C, Rohm H, Jaros D. How capsular exopolysaccharides affect cell surface properties of lactic acid bacteria. Microorganisms. 2020;8. doi:10.3390/microorganisms8121904

34. Barendt SM, Land AD, Sham L-T, Ng W-L, Tsui H-CT, Arnold RJ, et al. Influences of capsule on cell shape and chain formation of wild-type and pcsB mutants of serotype 2 Streptococcus pneumoniae. J Bacteriol. 2009;191: 3024–3040.

35. Gifford I, Suárez GA, Barrick JE. Evolution recovers the fitness of Acinetobacter baylyi strains with large deletions through mutations in deletion-specific targets and global post-transcriptional regulators. PLoS Genet. 2024;20: e1011306.

36. Renda BA, Dasgupta A, Leon D, Barrick JE. Genome instability mediates the loss of key traits by Acinetobacter baylyi ADP1 during laboratory evolution. J Bacteriol. 2015;197: 872–881.

37. Schembri MA, Dalsgaard D, Klemm P. Capsule shields the function of short bacterial adhesins. J Bacteriol. 2004;186. doi:10.1128/jb.186.5.1249-1257.2004

38. Kannisto M, Efimova E, Karp M, Santala V. Growth and wax ester production of an Acinetobacter baylyi ADP1 mutant deficient in exopolysaccharide capsule synthesis. J Ind Microbiol Biotechnol. 2017;44: 99–105.

39. Cossey SM, Velicer GJ, Yu YN. Ribonuclease D processes a small RNA regulator of multicellular development in myxobacteria. Genes (Basel). 2023;14. doi:10.3390/genes14051061

40. Romeo T. Global regulation by the small RNA-binding protein CsrA and the non-coding RNA molecule CsrB. Mol Microbiol. 1998;29: 1321–1330.

41. Bacher JM, Metzgar D, de Crécy-Lagard V. Rapid evolution of diminished transformability in Acinetobacter baylyi. J Bacteriol. 2006;188: 8534–8542.

42. Renda BA, Chan C, Parent KN, Barrick JE. Emergence of a competence-reducing filamentous phage from the genome of Acinetobacter baylyi ADP1. J Bacteriol. 2016;198: 3209–3219.

43. Liu B, Eydallin G, Maharjan RP, Feng L, Wang L, Ferenci T. Natural Escherichia coli isolates rapidly acquire genetic changes upon laboratory domestication. Microbiology. 2017;163: 22–30.

44. de Berardinis V, Vallenet D, Castelli V, Besnard M, Pinet A, Cruaud C, et al. A complete collection of single-gene deletion mutants of Acinetobacter baylyi ADP1. Mol Syst Biol. 2008;4: 174.

45. Wick RR, Judd LM, Cerdeira LT, Hawkey J, Méric G, Vezina B, et al. Trycycler: consensus long-read assemblies for bacterial genomes. Genome Biol. 2021;22: 266.

46. Kolmogorov M, Yuan J, Lin Y, Pevzner PA. Assembly of long, error-prone reads using repeat graphs. Nat Biotechnol. 2019;37: 540–546.

47. Wick RR, Holt KE. Polypolish: Short-read polishing of long-read bacterial genome assemblies. PLoS Comput Biol. 2022;18: e1009802.

48. Deatherage DE, Barrick JE. Identification of mutations in laboratory-evolved microbes from next-generation sequencing data using breseq. Methods Mol Biol. 2014;1151: 165–188.

49. Schindelin J, Arganda-Carreras I, Frise E, Kaynig V, Longair M, Pietzsch T, et al. Fiji: an open-source platform for biological-image analysis. Nat Methods. 2012;9: 676–682.

